# The Evolutionary History of the Orexin/Allatotropin GPCR Family: From Placozoa and Cnidaria to Vertebrata

**DOI:** 10.1101/403709

**Authors:** Alzugaray María Eugenia, Bruno María Cecilia, Villalobos Sambucaro María José, Ronderos Jorge Rafael

## Abstract

Cell-cell communication is a basic principle in all organisms, necessary to facilitate the coordination and integration between cell populations. These systems act by mean of chemical messengers. Peptides constitute a highly diversified group of intercellular messengers widely distributed in nature, and regulate a great number of physiological processes in Metazoa. Being crucial for life, it would seem that they have appeared in the ancestral group from which Metazoa evolved, and were highly conserved along the evolutionary process. Peptides act mainly through G-protein coupled receptors (GPCRs), a great family of transmembrane molecules. GPCRs are also widely distributed in nature being present not only in metazoan, but also in Choanoflagellata (unicellular eukariotes related with metazoans), and even in Fungi. Among GPCRs, the Allatotropin/Orexin (AT/Ox) family is particularly characterized by the presence of the DR**W** motif in the second intracellular loop (IC Loop 2), and seems to be present in Cnidaria, Placozoa and in Bilateria, suggesting that it also was present in the common ancestor of Metazoa. Looking for the evolutionary history of this GPCR family we searched in the GenBank for sequences corresponding to this family of receptors (i.e. seven transmembrane domain and the E/DRW motif at the second IC Loop 2). Our results show that AT/Ox receptors were highly conserved along evolutionary history of Metazoa, and that they might be defined by the presence of the E/DRWYAI motif at the level of IC Loop 2. Molecular phylogenetic analyses performed by Maximum Likelihood method suggest that AT/Ox family of receptors reflects evolutionary relationships that agree with current understanding of phylogenetic relationships in Actinopterygii and Sauropsida, including also the largely discussed position of Testudines.

## INTRODUCTION

Cell-cell communication is a basic principle in all organisms, necessary to facilitate the coordination and integration between cell populations, and with their environment. Indeed, integrative mechanisms as nervous and endocrine systems have appeared early along the evolutionary process and play a very important role, regulating many physiological processes in all animal phyla. As it is known, these systems act by mean of messengers which can be basically grouped as hormones and neuromodulators. Among these chemical messengers, peptides constitute a highly diversified group of molecules widely distributed in nature, and regulate a great number of physiological processes in most groups of Metazoa, from cardiac and visceral muscle activity, to more complex phenomena as sleep-wakefulness, and appetite.

Being this family of messengers crucial for life, it would seem that they have appeared in the ancestral group from which Metazoa evolved, and became highly conserved along the evolutionary process. Indeed, peptidic messengers are present in *Hydra sp*. and others members of the phylum Cnidaria [1–4], as well as in *Trichoplax adhaerens*, a member of the neuron-less animal phylum Placozoa [5–7], that also shares a common ancestor with Bilateria.

Peptides act mainly through G-protein coupled receptors (GPCRs), a complex and ubiquitous family of transmembrane molecules. GPCRs are widely distributed in Vertebrata, but also, this family of proteins, have been proved to be present in all metazoan, including Placozoa, Cnidaria, Ctenophora and Porifera, which share a common ancestor with Bilateria; also in Choanoflagellata (a group of unicellular eukariotes related with metazoans), and even in Fungi [1–3; 8–11].

GPCRs are characterized by the presence of seven transmembrane (TM) domains, an extracellular N-terminal and an intracellular C-terminal domains. The transmembrane domains are linked by three extracellular and three intracellular loops [for a review see 12, 13]. GPCRs are usually grouped in five major families, named *Rhodopsin, Frizzled, Glutamate, Adhesion* and *Secretin* [14]. Among these, the *Rhodopsin* family seems to be the most widely distributed in Metazoa and it is particularly characterized by the existence of a **<u>E</u>**/**<u>D</u>**R motif associated to the third transmembrane domain (TM3) (i.e. IC Loop 2), which seems to be relevant for the transmission of the message, facilitating the activity of the associated G-proteins [13, 14].

A vast number of the *Rhodopsin* family of receptor presents, as a conserved feature, the **E/D**R**Y/F** motif [14, 15]. In spite of that, a more limited number show the presence of a Tryptophan (W) instead that a Tyrosine (Y) residue (i.e. **E/D**RW). Among these, we found the receptors corresponding to the Allatotropin (AT) family of peptides [16].

AT is a neuropeptide originally isolated and characterized in insects on the basis of its ability to modulate the synthesis of Juvenile Hormones (JHs) in the gland corpora allata (CA) of the moth *Manduca sexta* (Lepidoptera: Insecta) [17]; and some other holometabolous species like the mosquito *Aedes aegypti* [18, 19]. Beyond the first biological function assigned, AT has proved to have multiple functions, including modulation of digestive enzymes secretion, and ion exchange regulation in the digestive system of Lepidotera [20, 21]. As a pleiotropic peptide, AT has also shown to be involved in myoregulatory processes, stimulating foregut movements in Lepidoptera [22]; and of the hindgut and midgut of both Chagas’ disease vectors *Triatoma infestans* and *Rhodnius prolixus* (Insecta: Hemiptera) [23, 24]. Furthermore, AT has proved to have cardioacceleratory functions synergizing the activity of serotonin in these species [24, 25]. In spite that AT was originally characterized as a neuropeptide (i.e. secreted by neurons at the central nervous system), it is also secreted by epithelial cells of the Malpighian tubules, and open-type cells at the level of the digestive system, acting in a paracrine and also endocrine way [25–28].

Looking for the evolutionary origin of allatoregulatory peptides, Alzugaray et al. [1, 2] have suggested that the AT/Ox and AST-C/somatostatin signaling systems are present in *Hydra sp*., a fresh water member of the phylum Cnidaria, playing myoregulatory roles during feeding, and modulating cytosolic Ca^2+^ levels [3]. Indeed, it was suggested that the allatotropic function of this peptides would constitute an insect synapomorphy, and that the ancestral function of these peptides could be myoregulatory [1, 29–31].

On the basis of a transcriptomic analysis performed in the CA/corpora cardiaca complex of the silkworm *Bombyx mori* the AT receptor (ATr) was identified [32]. Afterward, the receptor of AT in *M. sexta* was also characterized [33]. These authors confirmed that the receptor pertains to the *Rhodopsin* family of GPCRs, sharing a 48% of identity with the orexin receptor of vertebrates in the region comprised between the TM1 and TM7 domains [33]. Moreover ATr shares with orexin receptors the characteristic DR**W** motif [34].

Orexins (Ox), also named Hypocretins [35], originally identified in neurons located at the level of the hypothalamus in the rat, are two peptides sharing structural characteristics, derived from a same precursor by proteolytic processing [34, 35]. Initially related with physiological mechanisms regulating feeding behavior, the activity of these peptides was posteriorly associated with mechanisms regulating wakefulness and sleep [for a review see 36], and also with peripheral tissues activities. In fact, the presence of Ox and their receptors in the enteric nervous system, as well as at the level of the mucosa and smooth muscle of the digestive tract of mammals was also shown, suggesting that they also act as myoregulators [37, 38].

AT and Ox peptides are structurally different. Interestingly, bioinformatic search doesn´t show the presence of Ox in protostomates as well as AT in Deuterostomata, being possibly that, beyond the similarity between both receptors, Ox has evolved only in Deuterostomata and AT in Protostomata [1, 29, 30]. In fact, due that homology-based searches are often not sensitive enough to detect precursors of small peptides [5] and the difficulties to look for orthologues at the level of peptides, homologies between signal systems some times are based on their receptors [1; 39].

Looking for the evolutionary history of these signaling systems, we decide to go deeper in the analysis of these families of GPCRs (i.e. AT and Ox receptors). Based on fully characterized receptors both in vertebrates as well as in insects, we looked at the GenBank for putative AT/Ox receptors in all metazoan phyla. We have found sequences that might be considered AT/Ox GPCRs in several phyla including, Placozoa, Cnidaria, Mollusca, and Brachiopoda. On the basis of multiple sequence alignment we found motifs that might be considered “signatures” of the AT/Ox family of GPCRs. Phylogenetic analysis suggested that these families of receptors would be present in the ancestor of Metazoa, and that the system was highly conserved along evolutionary process. Moreover, a detailed maximum likelihood (ML) analysis of groups like Actinopterygii and Sauropsida, reflects phylogenetic trees that agree with current understanding of their phylogenetic relationships, including also the largely discussed evolutionary position of Testudines.

## 1 MATERIAL AND METHODS

### 2.1 Data retrieval

Sequences corresponding to Vertebrate and Insecta AT/Ox GPCRs were searched in protein database of the National Center for Biotechnology Information (NCBI) at https://www.ncbi.nlm.nih.gov/pubmed, and by protein BLAST (https://blast.ncbi.nlm.nih.gov/Blast.cgi?PROGRAM=blastp&PAGE_TYPE=BlastSearch&LINK_LOC=blasthome) on the basis of already annotated sequences in the Non-redundant protein sequences database. All the selected sequences were checked for the presence of the characteristic seven transmembrane domains using the TMHMM Server v. 2.0 (http://www.cbs.dtu.dk/services/TMHMM/). The presence of the **E**/**D**RW domain at the IC Loop 2 associated to TM3 was also verified. The sequences were then aligned using the Clustal Omega algorithm for multiple sequence alignment (http://www.ebi.ac.uk/Tools/msa/clustalo/) and further analyzed by the JalView 2.7 [40]. Only those sequences presenting the seven TMs and the **E**/**D**R**W** domains, were included.

### 2.2 Sequence analysis and alignment

Based on the alignment of the full set of sequences a search for motifs that might be considered as signatures in the AT/Ox family was performed. Once established at least one probable signature a search in different phyla including Bilateria and non-bilateria groups as Cnidaria y Placozoa were done. Each sequence were analyzed looking for both, the presence of the seven transmembrane domain pattern and the presence of the **E**/**D**R**W** motif. The phyla in which probable GPCRs associated to the AT/Ox family were found are:

Placozoa, Cnidaria, Arthropoda, Mollusca, Annelida, Brachiopoda and Chordata (see Supporting Information File 1).

### 2.3 Phylogenetic Analysis

Finally, the analysis of evolutionary relationships between sequences, except for the one corresponding to Fig. 1 (Neighbor-Joining), was performed using the ML method based on the Poisson correction model, including a 1000 replicates bootstrap analysis, with a 50% cut-off for condensed tree by the use of Mega 6.06 software [41]. The trees were then edited by the use of FigTree software (http://tree.bio.ed.ac.uk/software/figtree/).

**Figure 1.**
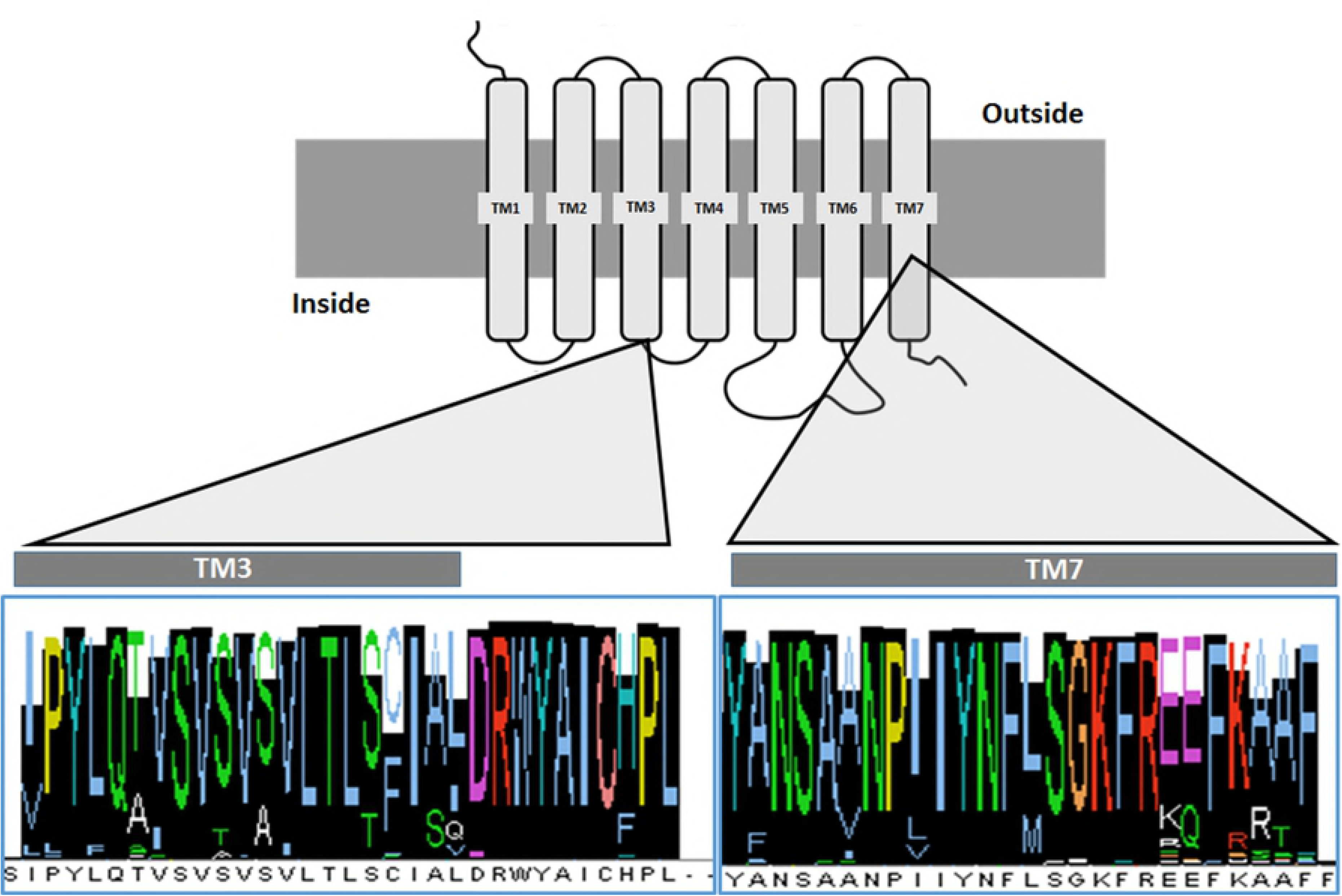
Schematic view of a generalized GPCR showing the two highly conserved domains, and the corresponding consensus after a multiple sequence alignment of sequences pertaining to the Allatotropin/orexin family of receptors. The alignment includes species pertaining to Placozoa, Cnidaria, Arthropoda, Mollusca, Annelida, Brachiopoda and Chordata.

The basic evolutionary relationships between groups are referred to *Tree of Life web Project* (http://tolweb.org/tree/) [42].

### 2.4 Search for signatures

Once the alignments were performed, we look manually for conserved motifs in different groups. The putative signatures were then blasted (https://blast.ncbi.nlm.nih.gov/Blast.cgi). Only those sequences presenting motifs covering the total length of the query blasted, showing %100 of identity were selected as putative signatures.

## RESULTS

### 3.1 The Allatotropin/Orexin receptors ancestral signature

As it is described above, GPCRs are characterized by the presence of the **E/D**R motif associated to the TM3 (i.e. IC Loop 2). Based on fully characterized AT and Ox receptors we looked in the GenBank for sequences in all animal phyla. After the analysis of 392 complete sequences, including N-terminal, C-terminal and the presence of 7 TM domains, we found that the motif **E/D**R**W**YI in the IC Loop 2 can be tracked from Chordata and Arthropoda, to Cnidaria and Placozoa. The most frequent motif found is **DRWYAI**, being present in 374 sequences, including the ancestral species *Trichoplax adhaerens* (Placozoa) (Table 1; Supporting Information File 1). The analysis of the rest of the sequences (eighteen), shows that seven of them exhibit only one conservative change, presenting ERWYAI corresponding to sequences of phyla pertaining to Lophotrochozoa (i.e. Mollusca, Brachiopoda and Annelida). The comparison of the codons codifying for the asparctic acid (D) and glutamic acid (E), shows that a point mutation at the third position of the codon would be responsible of this conservative change. A particular situation is presented in *H. vulgaris* (Cnidaria: Hydrozoa) in which the Tyrosine (Y) residue is substituted by asparagine (N), being the only sequence analyzed showing this conformation (i.e. ERW**N**AI). A point mutation at the first position of the codon should be responsible, and it has previously proposed as a sequence artifact [3].

**Table 1.** Characteristic Allatotropin/Orexin signature located at the interphase between transmembrane domain 3 (TM3) and the second intracellular loop (IC loop 2) distribution for every phylum analyzed.

| Phylum | Signature |
| --- | --- |
| Placozoa | **D/ERWYAI/V** |
| Cnidaria | **ERWYAI/V** |
| Brachiopoda | **ERWYAI** |
| Annelida | **ERWYAI** |
| Mollusca | **ERWYAI** |
| Arthropoda | **DRWYAI** |
| Chordata | **DRWYAI** |

### 3.2 Predicted sequences and general relationships between the animal phyla

As a result of a multiple sequence alignment, it also seems clear that at least two region of the AT/Ox receptor were highly conserved. One comprising the third transmembrane domain and its associated intracellular loop, and the second one comprising the TM7 (Fig. 1).

As a first approach to understand the relationships between the total sequences analyzed, a Neighbor-Joining analysis were performed (Fig. 2). The analysis shows that, as might be predicted, Placozoa (two sequences) and Cnidaria (three sequences pertaining to two different species of Anthozoa), clusters together sharing a common ancestor. Interestingly, the only sequence fitting the characteristics of the AT/Ox family of GPCR in *Hydra vulgaris* (Cnidaria: Hydrozoa) is clustered alone as the sister group of Bilateria (Fig. 2).

**Figure 2.**
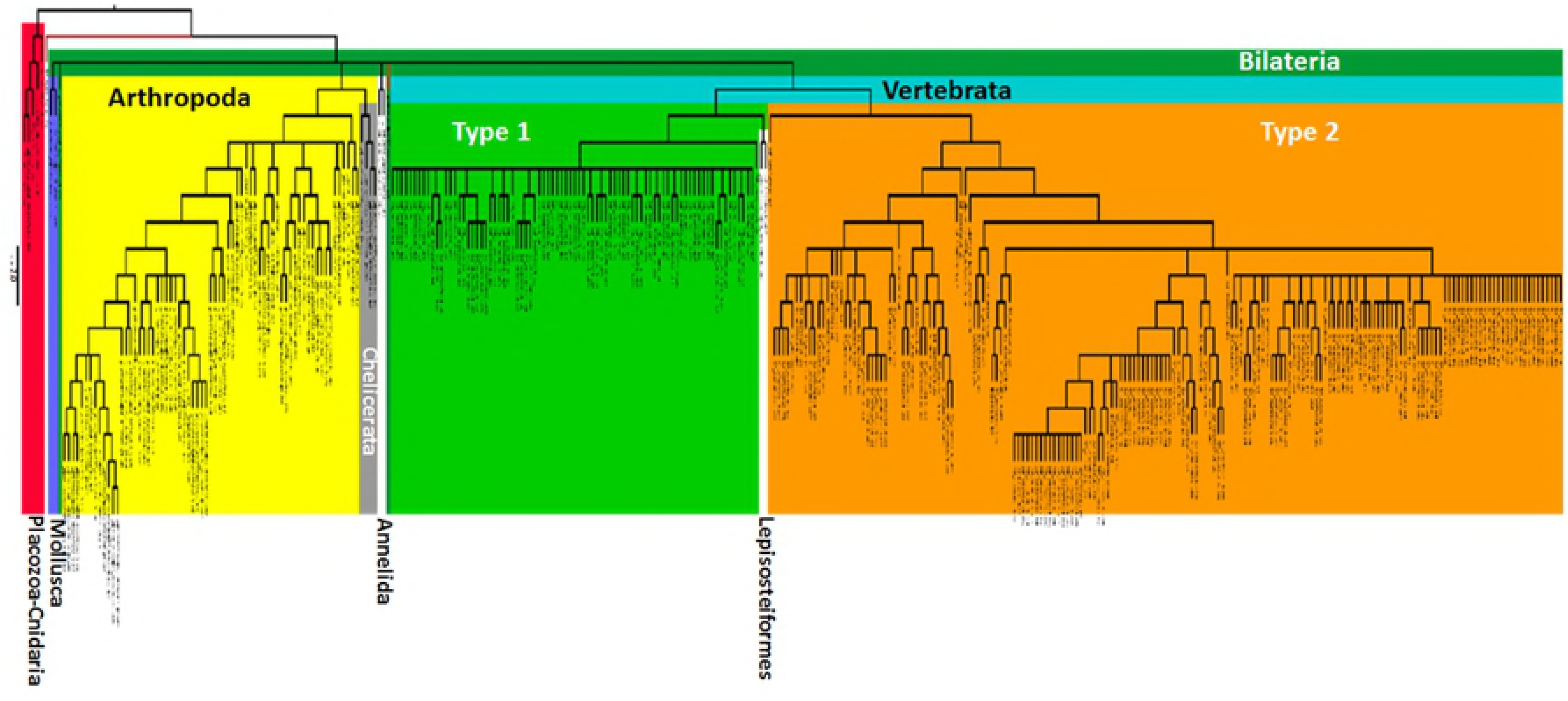
Evolutionary history of the Allatotropin/orexin family of receptors. All the sequences included present the seven transmembrane domains and the corresponding N-terminal and C-terminal domains. The tree was inferred by the Neighbor-Joining method. The cut-off value of replicate trees in which the associated taxa clustered together after a bootstrap test (1000 replicates) was 50%.

Despite of several genomes of the phylum Nematoda are fully sequenced, none of the GPCR sequences found in the GenBank showed the **DRWY** motif, suggesting that the AT/Ox system is not present in this phylum. Similar situation was found for the other two groups of Metazoa with uncertain positions as Porifera and Ctenophora.

Mammals is the only group of organisms in which the existence of two different kind of receptors was proved (i.e. Type 1 and Type 2), suggesting that the presence of these two receptors constitutes a synapomorphy of this group. Interestingly, *Lepisosteus oculatus*, pertaining to the group of Lepisosteiformes (with only six extant species), representing together with Halecostomi, the extant groups of Neopterygii, also presents two sequences, sharing the same clade with Type 1 receptor of Mammals. A more detailed analysis performed with ML methodology (see Fig. 3) also shows that these two sequences fit in the same clade, suggesting that the Type 1 Ox receptor appeared at least twice along the evolutionary history of Vertebrata (Fig. 2).

**Figure 3.**
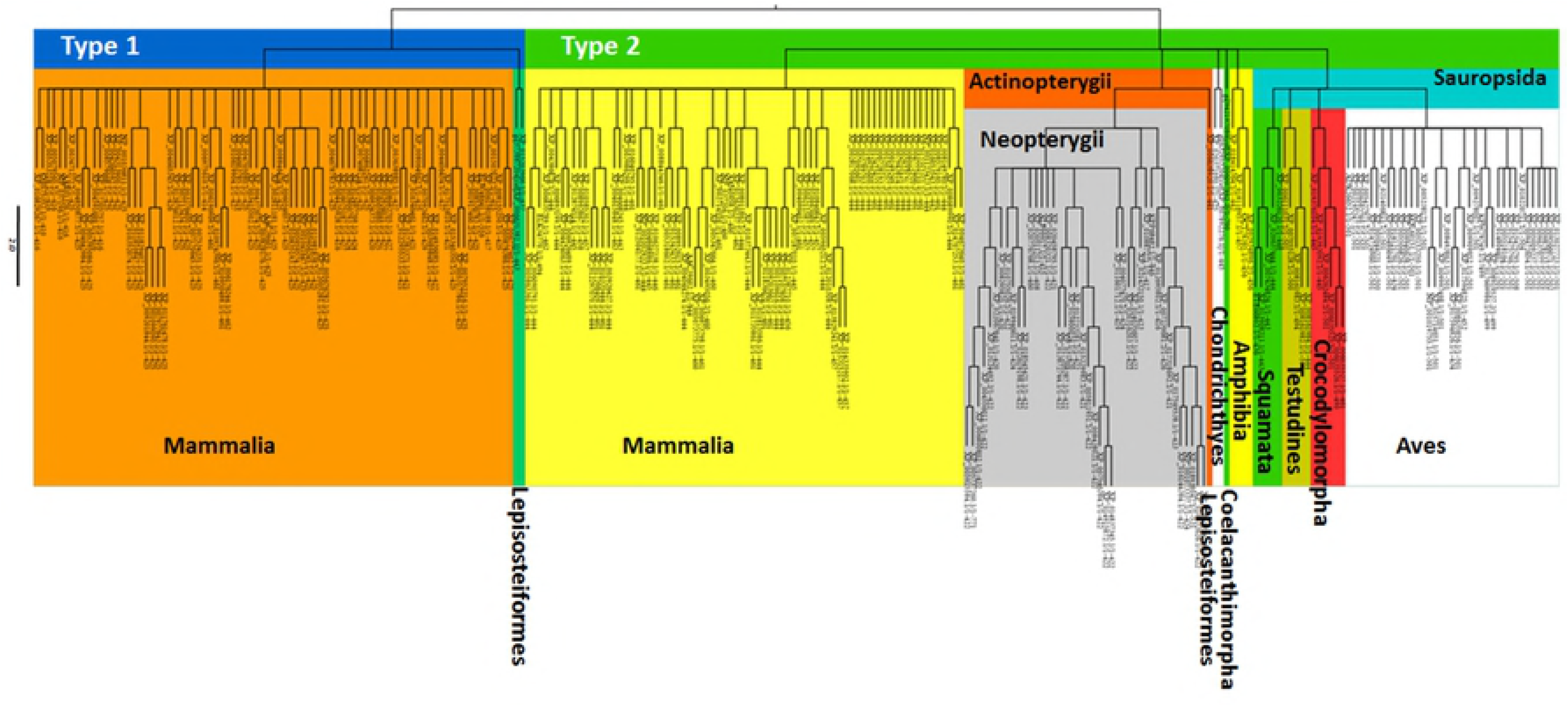
Phylogenetic relationships of Vertebrata. The tree was inferred by the Maximum Likelihood method. The cut-off value of replicate trees in which the associated taxa clustered together after a bootstrap test (1000 replicates) was 50%. Note that both kind of orexin receptors (Type 1 and Type 2) group independently. Type 2 receptor is present in all the groups of vertebrates included in the analysis. Type 1 receptor is only present in mammals with the exception of Lepisosteiformes (Actinopteygii: Neopterygii), suggesting that this kind of receptor could have appeared more than once along the evolution of Vertebrata.

Finally, the three best represented groups (i.e. Arthropoda, and the Type 1 and 2 receptors of Vertebrata) can be recognized at least by a highly conserved motif at the level of the interphase between TM3 and the second intercellular loop (see Table 2 and Supporting Information File 1).

**Table 2.**
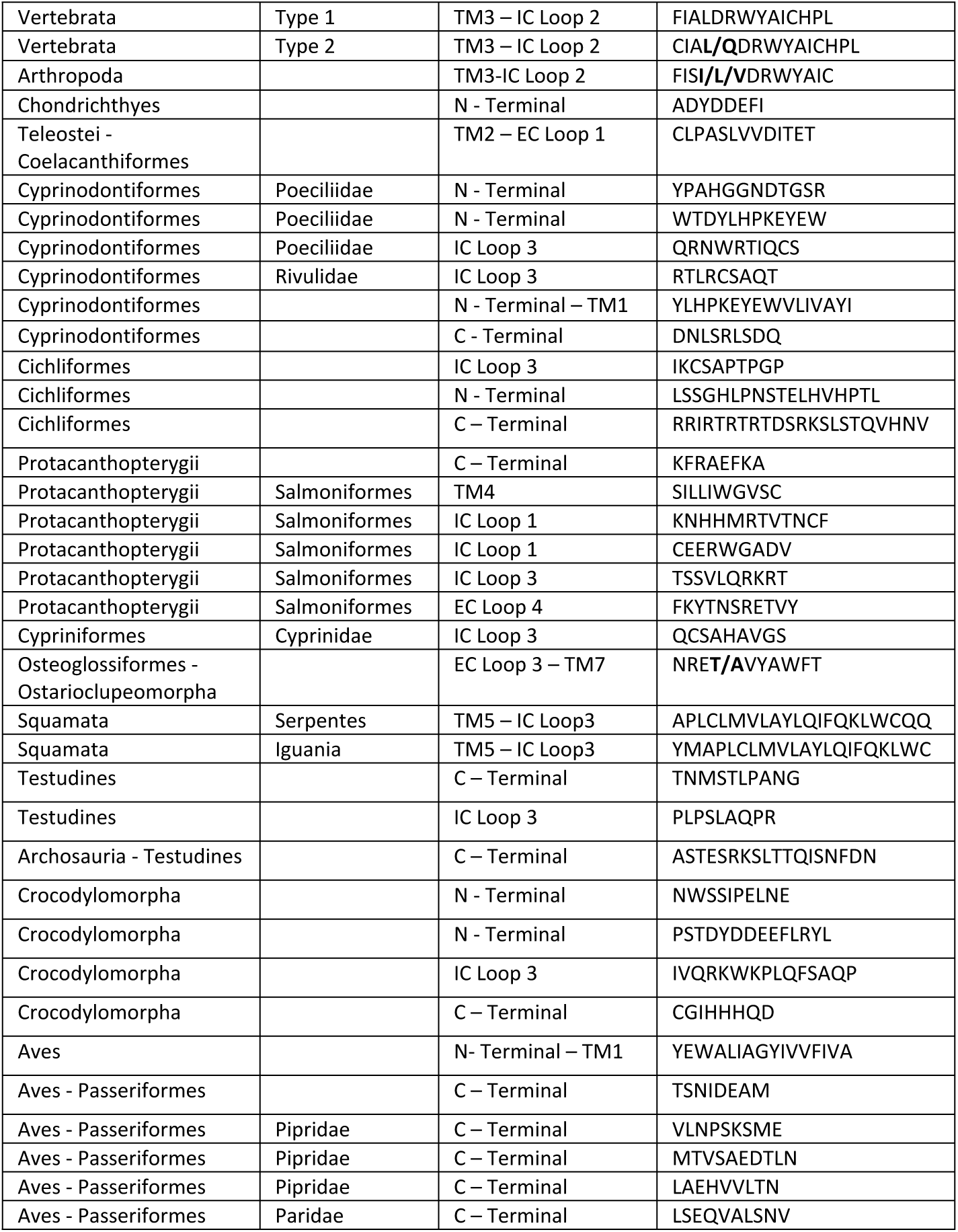
Putative signatures motifs and their location along the primary structure of the protein for AT/Ox GPCRs in different taxonomic groups. Note that most of the signatures are located at the level of the C-terminal domain (29.7%) and the N-terminal domain (21.6%).

### 3.3 Evolutionary history of orexin receptors in vertebrates

As previously stated, there exist two types of receptors in Vertebrate (i.e. Type 1 and Type 2). A ML analysis clearly divides the groups analyzed in two clades based on the Type 1 and Type 2 characteristics (Fig. 3). As we described above, two out of three sequences predicted for *L. oculatus* are grouped in the same clade of Type 1 receptor of Mammals. The other one (accession number XP_006638920) is grouped as a Type 2 receptor in the Actinopterygii clade (Fig. 3).

Regarding the Type 2 group, beyond that the Sarcopterygii are not grouped as a clade, showing Coelacanthimorpha, Amphibia, and the rest of tetrapoda a common ancestor with Actinopterygii and Chondrichthyes, the more represented groups (i.e. Mammals, Acitnopterygii and Sauropsida) are well defined as monophyletic groups (Fig. 3).

### 3.4 Sauropsida

As a first attempt to further understand the evolutionary history of the Ox receptor family, we decide to go deeper in the analysis of two groups of vertebrates well represented in our sample, as Sauropsida and Actinopterygii are, looking also for signatures motifs for every group analyzed. In fact, after a detailed analysis of the alignments for each group, we could find signature motifs, that once blasted in the GenBank, remitted specifically to most of the groups under study (Table 2).

A ML analysis of Sauropsida shows two well supported clades conformed, one of them by Lepidosauromorpha species, including those corresponding to Iguania and Serpentes, traditionally grouped in the order Squamata, and the second one, conformed by Archosauria and Testudines (Fig. 4). Regarding Squamata, sequences in the TM5 – IC Loop 3 seems to be characteristic, showing Serpentes and Iguania the motifs **APLCLMVLAYLQIFQKLWCQQ** and **MAPLCLMVLAYLQIFQKLWC** respectively (Table 2).

**Figure 4.**
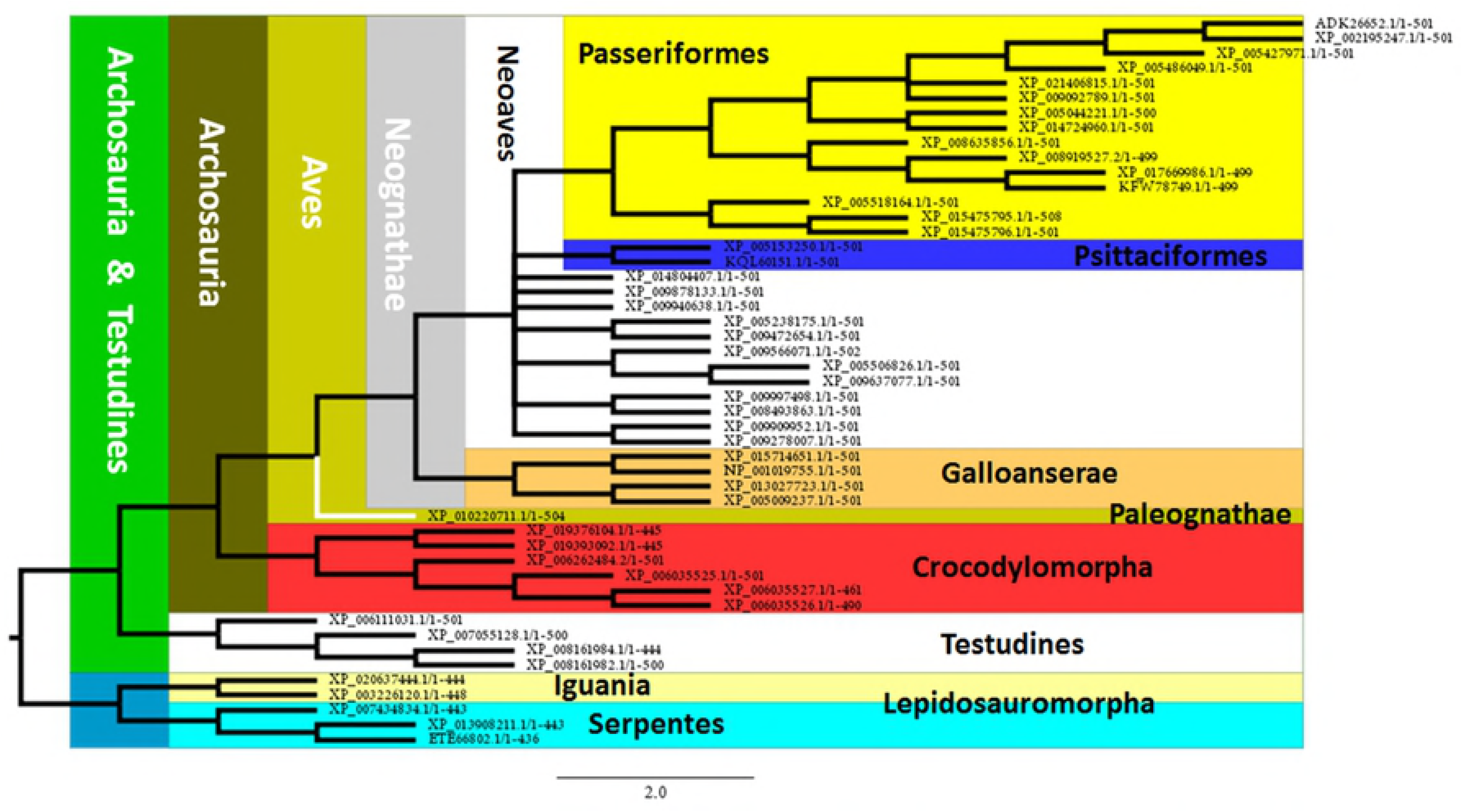
Maximum Likelihood analysis of Sauropsida. The phylogeny is clearly represented showing Lepidosauromorpha as the sister group of Archosauria. The main groups of Aves are also represented. Two orders of Neoaves (Passeriformes and Psittasiformes) are recognized. Furthermore, in Passeriformes, the best represented group, two families can be recognized by signature motifs. Testudines appear as the sister group of Archosauria in agreement with the current accepted hypothesis that recognize them as Diapsida, resembling also the currently proposed group of Archelosauria.

As would be expected, Archosauria presents a well-defined phylogenetic pattern involving Crocodylomorpha, with species representing the three extant groups (i.e. Gavialoidea; Alligatoroidea and Crocodyloidea) and Aves. The clade including Crocodylomorpha seems to be characterized for four different signature motifs; two located at the N-terminal domain, one corresponding to the C-terminal, and a third one in the IC Loop 3 (see Table 2). With respect to the birds, a sequence located in the interphase between N-terminal and TM1 would act as a signature (Table 2).

The clade corresponding to Aves, currently accepted as members of Coelurosauria (Dinosauria: Saurischia), shows the sequence **YEWALIAGYIVVFIVA** in the interphase N-terminal – TM1, fully conserved (Table 2). With respect to the phylogenetic relationships, the main groups are represented and grouped as well, including Paleognathae (*Tinamus guttatus*), and Neognathae which in fact form two well supported clades including Galloanserae and Neoaves (Fig. 4).

Moreover, the two groups of Galloanserae are represented by four species pertaining to different genus, grouped in the expected clades. In fact, *Anser cygnoides* and *Anas platythynchos* (Anseriformes), and *Coturnix japonica* and *Gallus gallus* (Galliformes) form two monophyletic groups. Regarding the Neoaves, only two currently recognized orders, Psittaciformes (represented by two species) and Passeriformes, are well defined (Fig. 4). Passeriformes represented by 16 sequences, would be recognized by the sequence **TSNIDEAM** at the C-terminal domain. Moreover, two families in this group, Pipridae and Paridae, would also be identified by signatures at the level of the C-terminal domain (Table 2).

The last point to analyze is the position of turtles which phylogenetic position have been largely discussed. Our analyses shows the clade of Testudines, represented by species pertaining to three different families, as the sister group of Archosauria (Crocodylomorpha + Aves). Indeed, the sequence **ASTESRKSLTTQISNFDN** corresponding to the C-terminal domain, identify the Archosauria-Testudines clade (Fig. 4, Table 2).

### 3.5 Actinopterygii

Regarding to Actinopterygii (represented by species corresponding only to Neopterygii), the ML analyses of Type 2-like receptor, present them as a well-supported clade, sharing a common ancestor with Condrichthyes which are characterized by the presence of the **ADYDDEFI** motif at the level of the N-terminal (Fig. 5, Table 2). As expected, the sequence corresponding to Type 2 receptor of Lepisosteiformes appears as the sister group of Halecostomi (Fig. 5). With respect to Halecostomi, only sequences corresponding to Teleostei was found. Amiiformes, one of the extant group is not represented in our samples. Teleostei, the more diversified group, represented by numerous species that can be grouped in 11 different clades (see tolweb.org for reference) is represented by 6, including Osteoglossomorpha, Ostariophysi, Clupeomorpha, Salmoniformes, Esociformes and Acanthomorpha (Fig. 5). Similarly to other studies, Osteoglossomorpha (Scleropages formosus) appears as the sister group of the clade that includes Ostarioclupeomorpha (Ostariophysi and Clupeomorpha) and Euteleostei (Protacanthopterygii and Neoteleostei).

**Figure 5.**
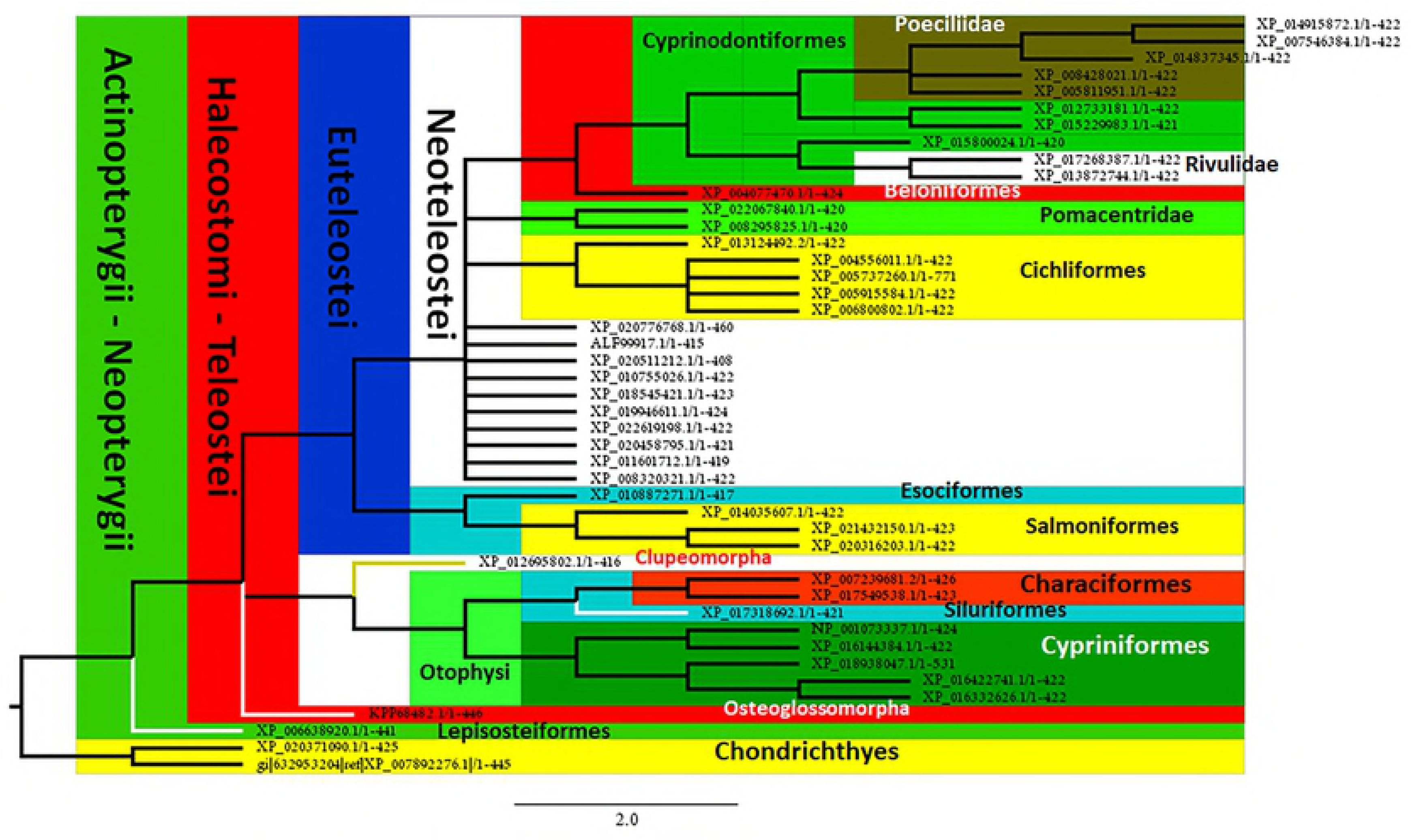
Analysis of Maximum Likelihood of sequences of Orexin receptor corresponding to Actinopterygii. All the species pertain to Neopterygii being represented the two extant groups (Lepisosteiformes and Halecostomi), which appear as the sister group of Chondrichthyes. Currently proposed groups are clearly represented at higher taxonomic levels. The analysis also recognizes taxa at lower levels including families defined by characteristic motifs that might be considered as signatures.

The other two clades of Teleostei (i.e. Ostarioclupeomorpha and Euteleostei) share a common ancestor. The first one, involves one Clupeomorpha species appearing as the sister group of Otophysi, which is well represented by three out of four recognized orders (Characiformes, Siluriformes and Cypriniformes) (Fig. 5). Indeed, Characiformes and Siluriformes are grouped in a clade as expected by previous phylogenetic studies, being the sister group of Cypriniformes. Regarding Euteleostei, the two main clades appear as sister groups; Protacanthopterygii (which could be characterized by the presence of the **KFRAEFKA** motif in the C-terminal), including Esociformes (*Esox lucius*) and Salmoniformes. Salmoniformes are represented by three species of two different genus: *Salmo salar, Oncorhynchus mykiss* and *O. kisutch*. Moreover, the two species of the genus *Oncorhynchus* are recognized as a clade (Fig. 5). Regarding Salmoniformes, our analyses show the existence of 5 different motifs that might be considered as signatures (see Table 2).

With respect to Neoteleostei, a total of 28 sequences were analyzed, pertaining all of them to the clade of Percomorpha (Acanthopterygii), corresponding to: Pleuronectiformes (2), Gasterosteiformes (1), Synbranchiformes (1), Tetraodontiformes (1), Beloniformes (1), Cyprinodontiformes (10) and 12 species corresponding to the non-monophyletic traditional “Perciformes”. The members of two families traditionally considered as members of the order Perciformes, as Pomacentridae (represented by two species) and Cichlidae (five species), are well grouped as individual clades. Indeed, the clade of Cichlidae, currently considered as the Order Cichliformes [43] might be identified by three different motifs located at the N-terminal, C-terminal, and IC Loop 3 (Fig. 5, Table 2). Finally, other well represented group is Cyprinodontiformes, characterized by the presence of **DNLSRLSDQ** motif at the C-terminal domain, including 10 sequences corresponding to five different families, being Rivulidae (2) and Poeciliidae (5 species) those best represented. Interestingly both of them are grouped as individual clades (Fig. 5), being characterized by the **RTLRCSAQT** (Rivulidae) and **QRNWRTIQCS** motifs (Poeciliidae). Regarding Poeciliidae, two more motifs might be characteristics at the level of the N-terminal domain (Table 2).

## DISCUSSION

As it is known, GPCRs are widely distributed in nature, being associated with the regulation of a great number of physiological mechanisms. As they are engaged with critical processes it is not rare that they were conserved along the evolutionary processes, being appeared early in the evolution. Indeed, SWSI (short-wavelength sensitive opsin), another member of the GPCR family of proteins which is involved in light signal transduction, has proved to be a potential phylogenetic marker in Vertebrata, showing phylogenetic relationships congruent with the evolution of this group at both high and low taxonomic levels [44].

We have previously shown that GPCRs are present in a variety of Metazoa, including *T. adhaerens*, the multicellular organism pertaining to the neuron-less phylum, Placozoa [1]. Moreover, previous studies in our laboratory suggest that in *Hydra sp*. GPCRs associated with regulatory peptides are present (Cnidaria: Hydroazoa) [1–3]. Indeed, these studies suggest the existence of Allatotropin/Orexin and Allatostin-C/Somatostatin homologous systems that would act as myomodulators, controlling the movements associated with capture and digestion of the prey in *Hydra sp.* [1–3].

Regarding AT/Ox GPCRs, as we detailed above, they are characterized by the presence of a Tryptophan (W) instead of a Tyrosine (Y) associated to the E/DR motif in the IC Loop 2 [16]. Our results show, that the AT/Ox family of GPCRs may be defined by the presence of the **E/DRWYAI** motif, present in 381 out of 392 sequences analyzed, covering most of the Metazoa phyla, and that might be considered as a signature of the family. Furthermore, despite we could not find any convincing sequence showing this characteristic motif nor in Ctenophora neither in Porifera, due to its presence in Placozoa, Cnidaria and Bilateria, it might be assumed that the AT/Ox family of GPCRs was present in the common ancestor of Metazoa. The lack of the AT/Ox family of GPCR in those phyla, might be a biological phenomenon, or perhaps an artifact. In fact, beyond the great quantity of information about genomic and transcriptomic sequencing, it may be assumed that it is still perfectible [45]. Indeed, the phylogenetic positions and the evolutionary relationships between Ctenophora, Porifera and the rest of the metazoan groups is still controversial [8, 46]. Moreover, regarding GPCRs, it was already suggested that the Porifera Rhodopsin family has not orthologous relationship with the ones found in the rest of Metazoa [11].

Regarding Vertebrata two different groups were found. Interestingly, are not defined by their phylogenetic relationships, but by the kind of the protein constituting the receptor (Type 1 and Type 2 receptor). One of these groups (i.e. Type 2) is represented in all the groups including, Chondrichthyes, Actinopterygii, Sauropsida and Mammalia, and might be defined for the presence of the **CIAL/QDRWYAICHPL** motif. On the other hand, with the exception of Lepisosteiformes (Actinopterygii: Neopterygii), Type 1 receptor is exclusively expressed in Mammalia (defined by the **FIALDRWYAICHPL** motif). In fact, in Lepisosteiformes, three different sequences were found; two of them are grouped in all the analysis performed with the Type 1 receptor of mammals showing also the **FIALDRWYAICHPL** motif in the interphase between TM3 and the IC loop 2. Beyond these two sequences, a third one (grouped as Type 2 receptor), shows a phylogenetic position according to the current assumption, as the sister group of Halecostomi. The existence of two kind of Ox receptors might be considered as a synapomorphy of Mammalia. The presence of the Type 1-like receptor in Lepisosteiformes would be suggesting that this receptor had appeared more than once along the evolution of Vertebrata.

As a way to further understand the evolutionary history of this family of receptors, we decided to go deeper in the analysis of Type 2-like receptor phylogenetic relationships in two groups of Vertebrata (Sauropsida and Actinopterygii). In both of them, our results show that the sequences phylogenetic relationships are mostly in agreement with current hypothesis about their phylogeny. As an example, a group of species of Neoteleostei (i.e. *Oreochromis niloticus, Maylandia zebra, Neolamprologus brichardi, Haplochromis burtoni* and *Pundamilia nyererei*), traditionally considered as the Cichlidae family pertaining to the order Perciformes (at present considered as polyphyletic), are still grouped as a clade, that in fact is now considered as the order Cichliformes [43]. Another interesting point is that related with the order Cyprinodontiformes. This group represented by 10 species pertaining to five different families, are well defined as independent groups, being the two families represented by two or more species (i.e. Poeciliidae and Rivulidae) grouped as monophyletic groups sharing a common ancestor with the rest of the species of the order. Indeed, these two families might be recognized by signatures located at the N-terminal and IC Loop 3.

Other interesting subject is related with the phylogeny of Sauropsida and the evolutionary position of turtles (Testudines). The phylogenetic position of turtles was largely controversial, as they were traditionally considered as an order pertaining to the group of Anapsida (having no temporal fenestrae in their skull). Traditional studies based on paleontological and morphological characters positioned them as the only extant group of Anapsida being the sister group of Diapsida (a clade that includes Lepidosauromorpha, Archosauria as sister groups). Based on both paleonthologycal and molecular phylogeny, the evolutionary relationships of Testudines was revisited, considering them as the sister group of Lepidosauromorpha, or as the sister group of Archosauria (Aves and Crocodylomorpha) [for a review see 47]. The finding of a stem-turtle from the middle Triassic finally positioned turtles as a member of Diapsida [48, 49]. In agreement with previous molecular studies [50–53], our results, based on sequences pertaining to three different families, place Testudines as the sister group of Archosauria, sharing the **ASTESRKSLTTQISNFDN** motif at the C-terminal domain. Indeed the existence of a new group including Testudines and Archosauria named Archelosauria was recently proposed [51].

Finally, our results show the existence of numerous motifs that might be considered as signatures for several of the groups analyzed, being hypothetically possible to test them both as phylogenetical markers at both higher and lower taxonomic levels.

## ACKNOWLEDGMENTS

This research was supported by funds provided by the National University of La Plata (N/813) and CONICET (PIP N° 112201-50100419). MEA and MJVS are researcher of CONICET (Argentina). The authors also wish to thanks to Dr. Cecilia Morgan for the critical reading of the manuscript and her valuable comments.

## Supporting Information file 1

Complete list of sequences analyzed. The putative signatures motifs for every group are highlighted.

